# Cross-validation of technologies for genotyping *CYP2D6* and *CYP2C19*

**DOI:** 10.1101/2019.12.24.870295

**Authors:** Beatriz Carvalho Henriques, Amanda Buchner, Xiuying Hu, Vasyl Yavorskyy, Yabing Wang, Kristina Martens, Michael Carr, Bahareh Behroozi Asl, Joshua Hague, Wolfgang Maier, Mojca Z. Dernovsek, Neven Henigsberg, Daniel Souery, Annamaria Cattaneo, Joanna Hauser, Ole Mors, Marcella Rietschel, Gerald Pfeffer, Chad Bousman, Katherine J. Aitchison

## Abstract

**Background:** CYP2D6 and CYP2C19 are cytochrome P450 enzymes involved in the metabolism of many medications from multiple therapeutic classes. Associations between patterns of variants (known as haplotypes) in the genes encoding them (*CYP2D6* and *CYP2C19*) and enzyme activities are well described. The genes in fact comprise 21% of biomarkers in drug labels. Despite this, genotyping is not common, partly attributable to its challenging nature (*CYP2D6* having >100 haplotypes, including those with sequence from an adjacent pseudogene, and gene duplications). We cross-validated different methodologies for identifying haplotypes in these genes against each other.

**Methods:** Ninety-two samples with a variety of *CYP2D6* and *CYP2C19* genotypes according to prior AmpliChip CYP450 and TaqMan *CYP2C19*17* data were selected from the Genome-based therapeutic drugs for depression (GENDEP) study. Genotyping was performed with TaqMan copy number variant (CNV) and single nucleotide variant (SNV) analysis, the next generation sequencing-based Ion S5 AmpliSeq Pharmacogenomics Panel, PharmacoScan, long-range polymerase chain reaction (L-PCR) followed by amplicon analysis, and Agena for *CYP2C19*. Variant pattern to haplotype translation was automated.

**Results:** The inter-platform concordance for *CYP2C19* was high (up to 100% for available data). For *CYP2D6*, the IonS5-PharmacoScan concordance was 94% for a range of variants tested apart from those with at least one extra copy of a *CYP2D6* gene (occurring at a frequency of 3.8%, 33/853), or those with substantial sequence derived from pseudogene, known as hybrids (3%, 26/853).

**Conclusions:** Inter-platform concordance for *CYP2C19* was high, and, moreover, the Ion S5 and PharmacoScan data were 100% concordant with that from a TaqMan *CYP2C19*2* assay. We have also demonstrated feasibility of using an NGS platform for genotyping *CYP2D6* and *CYP2C19*, with automated data interpretation methodology. This points the way to a method of making *CYP2D6* and *CYP2C19* genotyping more readily accessible.

## Introduction

CYP2D6 and CYP2C19 are cytochrome P450 enzymes involved in hepatic phase I reactions that alter the hydrophobicity and reactivity of substrates. Substrates for CYP2D6 include at least 25% of commonly prescribed medications from a variety of classes, including antidepressants (e.g., venlafaxine, citalopram, fluoxetine, paroxetine, amitriptyline, nortriptyline, imipramine, desipramine, clomipramine, trimipramine) antipsychotics (e.g., aripiprazole, risperidone, haloperidol, zuclopenthixol, and pimozide), antiarrhythmics (e.g., flecainide and propafenone), beta-blockers (e.g., metoprolol and propranolol), opioids (e.g., codeine), antiemetics (e.g., metoclopramide and ondansetron), antitussives (e.g., dextromethorphan) and anticancer agents such as the hormonal modulator tamoxifen.^1–4^ Substrates for CYP2C19 include proton pump inhibitors (omeprazole and pantoprazole), the antithrombotic clopidogrel, antidepressants (fluoxetine, sertraline, paroxetine, citalopram, escitalopram, moclobemide, imipramine, amitriptyline, clomipramine), antipsychotics (e.g., clozapine and olanzapine), anxiolytics (diazepam and propranolol), the anticonvulsant phenytoin, and immunologically active agents (e.g., cyclophosphamide, nelfinavir, proguanil, and voriconazole).^4^

The gene (*CYP2D6*) encoding the enzyme CYP2D6 is on chromosome 22q13.2^5^ adjacent to two homologous pseudogenes, *CYP2D7* and *CYP2D8*.^6^ This arrangement is susceptible to unequal crossover events in meiosis. For example, *CYP2D6* pairs with *CYP2D7* or part thereof, resulting in a deletion of the *CYP2D6* gene on one chromatid and an extra copy of *CYP2D6* on the other. In addition, it has been postulated that other recombination mechanisms,^7^ such as those facilitated by the multiple transposable genetic elements flanking all three *CYP2D* genes,^8^ have contributed to locus variability. The transposable elements include long and short interspersed nuclear elements (the origin of which includes reverse-transcribed viral RNA). For example, repeat regions comprising such sequences between *CYP2D7* and *CYP2D6* and downstream of *CYP2D6* are thought to have facilitated mechanisms leading to the formation of hybrid genes made up of sequence derived in part from *CYP2D7* and in part from *CYP2D6*^7,9^. These various types of chromosomal rearrangements, otherwise known as structural variants, have been historically challenging to detect by many different types of genotyping techniques. In addition, the region is one of the most variable areas of the human genome, with over 100 different haplotypes or “star alleles” (defined by particular genetic variants or combinations thereof on the same chromosome) having been identified and catalogued.^10,11^ Moreover, new variants continue to be discovered. It is therefore hardly surprising that genotyping of *CYP2D6* for treatment optimization is not being routinely provided in clinical settings, despite the presence of guidelines indicating potential clinical utility.^12–14^

The *CYP2C19* gene encoding the CYP2C19 enzyme is likewise located at chromosome 10q23.33 together with other similar genes, namely *CYP2C18* (not expressed at the protein level), *CYP2C9*, and *CYP2C8*.^15–17^ While structural variants of *CYP2C19* have recently been identified,^18^ the more commonly studied haplotypes result from single base changes in the DNA sequence (single nucleotide variants or SNVs); more than 30 different *CYP2C19* haplotypes have been catalogued to date.^11^

Each *CYP2D6* or *CYP2C19* haplotype is associated with different levels of enzyme activity, ranging from haplotypes of loss-of-function (which give rise to no functional enzyme), to decreased function (which are associated with an enzyme with reduced metabolic activity), to gain-of-function (associated with increased activity).^2,4^ The CYP2D6 and CYP2C19 enzyme activity groups vary markedly by ethnic group. CYP2D6 UMs have been found at a frequency of 0.9 - 4% in Whites and at higher frequency in Sub-Saharan Africa; CYP2D6 IMs are relatively common in East Asia and in Blacks; and CYP2D6 PMs are relatively common in Caucasians at 7% and rare in East Asians at 1%.^1,19,20^ The frequency of CYP2C19 PMs is 2-5% in Caucasians, 2% in Saudi Arabians, 4% in Zimbabweans, 5% in Ethiopians, 13% in Koreans, 15-17% in Chinese, 21% in Indians, and 18-23% in Japanese.^1,4,21,22^

There is evidence that genotyping for CYP2D6 and CYP2C19 would reduce the number of adverse drug reactions (ADRs) to medications metabolized by these enzymes. For example, Pirmohamed et al. (2004)^23^ found that antidepressants were the leading mental health class of medication and fifth leading medication class overall of ADR-related general hospital admissions. Methodology for genotyping these genes effectively and efficiently could therefore lead to substantial health care costs savings. The first step in introducing a new genetic test to a clinical genetics laboratory is establishing analytical validity. Our aim was to contribute to this step by cross-validating multiple technologies for genotyping *CYP2D6* and *CYP2C19* against each other and providing automated translation of data with multiple variants into haplotypes.

## Methods

Ninety-two DNA samples originating from venous blood samples were selected from those previously genotyped for *CYP2D6* and *CYP2C19* using the AmpliChip CYP450 supplemented by the TaqMan assay C_469857_10 for *CYP2C19*17* as part of the Genome-based therapeutic drugs for depression (GENDEP) study24. Participants were all of self-reported White European ancestry. The AmpliChip identified 32 *CYP2D6* variant haplotypes (*\*2, *3, *4, *5, *6, *7, *8, *9, *10, *11, *14, *15, *17, *19, *20, *25, *26, *29, *30, *31, *35, *36, *40, *41, *1XN, *2XN, *10XN, *17XN, *35XN, *41XN*). In addition, it covered *CYP2C19* haplotypes *\*2* and *\*3*. The non-variant (known as wild-type) haplotype (*\*1*) for each gene was identified by default as an absence of the mutations defining the specific variants tested. Samples were quantified using fluorimetry-based methods (Qubit or Quantifluor).

### TaqMan copy number variant (CNV) analysis and sample selection

TaqMan copy number assays for *CYP2D6* (assay IDs Hs04083572_cn and Hs00010001_cn for intron 2 and exon 9 respectively, Thermo Fisher Scientific) were run according to the manufacturer’s protocol on a ViiA7 Real-Time PCR System (Thermo Fisher Scientific). The assays are specific for intron 2 and exon 9 in *CYP2D6*, with DNA not being amplified if the relevant region is *CYP2D7* sequence. Assays were run in quadruplicate with internal calibrators of known *CYP2D6* copy number. Data were analysed using CopyCaller software version 2.1 (Thermo Fisher Scientific) with internal calibrators of known *CYP2D6* copy number according to the manufacturer’s instructions (using a confidence level of at least 95%, most being above 99%).

Samples for which the TaqMan CNV call across the two probes were not equal were identified as putative hybrid haplotypes^25^ and analyzed with a third probe (assay ID Hs04502391_cn for *CYP2D6* intron 6). These samples and those with a unique combination of *CYP2D6* and *CYP2C19* genotypes by the AmpliChip CYP450 and TaqMan *CYP2C19*17* assays, or no reference sample for that genotypic combination available at the time of selection (June 2019), or a no call for *CYP2D6* on the AmpliChip CYP450 were then taken forward for further analysis. The following genotyping techniques were employed: PharmacoScan (Thermo Fisher Scientific), Ion S5 AmpliSeq Pharmacogenomics Panel (Thermo Fisher Scientific), and TaqMan Drug Metabolism Genotyping Assays (Thermo Fisher Scientific). Data arising from these were then used to select samples for the generation of amplicons by long-range polymerase chain reaction (known as L-PCR) for the detection of specific gene *CYP2D6* duplications and conversions (hybrids).^9,26,27^ In addition, a TaqMan assay for *CYP2C19*2* (assay ID C_25986767_70) was used as a further validation test for this SNV and Agena was used for *CYP2C19*.

### Ion S5 AmpliSeq Pharmacogenomics Panel

Genotyping using the Ion S5 AmpliSeq Pharmacogenomics Panel (Thermo Fisher Scientific) was conducted according to the manufacturer’s instructions using an Ion Chef instrument (Thermo Fisher Scientific, Waltham MA). Short stretches of genomic DNA were sequenced, including regions of *CYP2D6* designed to detect gene deletion, duplication and conversion (*CYP2D6-2D7* hybrid) events. In brief, library preparation was conducted using the Ion AmpliSeq Library Kit Plus (genomic DNA was subjected to a multiplex PCR using a pool of primer pairs). The resulting amplicons were then partially digested and ligated to barcode adaptors (Ion Express barcode adaptors 1-96 kit, Thermo Fisher Scientific), which were then purified using a magnetic bead technology (AMPure XP Reagent, Agencourt). Library quantitation was carried out by quantitative PCR using the Ion Library TaqMan Quantitation Kit (Thermo Fisher Scientific) and was followed by normalisation of library concentration to 40 pM. Two chips were then loaded, containing libraries 1-48 and 49-96 respectively. Following sequencing, data were accessed remotely using the GeneStudio Data Analysis software (Thermo Fisher Scientific). Ion S5 sequencing generated an average of 109,454 reads per sample (mean read length 142.5 bp), with two samples failing quality control (in a manner indicating likely insufficient template: mapped read numbers of 18 and 51). Variant calling by the Ion Torrent Variant Caller version 5.10.1.19 (Thermo Fisher Scientific) generated three text files: one with the genotype at each SNV (including 20 *CYP2D6* variants and 11 *CYP2C19* variants), one for the *CYP2D6* exon 9 CNV output, and one for the *CYP2D6* gene level CNV data (based on sequence across nine regions in *CYP2D6*). Haplotype translation files were created to derive *CYP2D6* and *CYP2C19* haplotypes including various hybrid configurations from the SNV and CNV data using the AlleleTyper software (Thermo Fisher Scientific). These translator files are available at request.

### PharmacoScan

PharmacoScan is an array-based technology in which the sample DNA is hybridized to short lengths of DNA pre-bound to a “gene chip;” this assay was run at Neogen Genomics (Nebraska, USA). Genomic DNA was amplified (including preferential amplification of *CYP2D6* and *CYP2C19*), with the amplicons then being fragmented, labelled, and hybridized to a PharmacoScan Array (Thermo Fisher Scientific). Arrays stained with a fluorescent antibody were scanned on a GeneTitan Multi-Channel Instrument (Thermo Fisher Scientific). The resultant data including more than 100 variants in *CYP2D6* and 60 variants in *CYP2C19* were analysed using the Axiom Analysis Suite 4.0.3.3 (Thermo Fisher Scientific). The latest version (v8.2) of the manufacturer’s haplotype translation file was used. CNV calls were provided by probes for exon 9 of *CYP2D6* as well as for the 5’ and 3’ flanking regions as described.^28^

### Long-range PCR assays with characterisation of resultant amplicons

Haplotype assignment for samples with a call of three across the TaqMan or Pharmacoscan CNV probes was conducted by L-PCR as described to generate an amplicon specific for the duplicated gene^27^ followed by gel extraction (GeneJET Gel Extraction Kit, Thermo Fisher Scientific, Waltham, MA) and the use of TaqMan Drug Metabolising assays specific for the haplotypes found in samples of heterozygous *CYP2D6XN* genotype by at least one platform (such as \**2*, *\*3*, *\*4*, *\*6*, *\*35* and *\*41;* TaqMan assay IDs C_27102425_10, C_32407232_50, C_27102431_D0, C_32407243_20, C_27102444_F0, and C_34816116_20, respectively). Samples were run in duplicate on a ViiA7 Real-Time PCR System (Thermo Fisher Scientific), using automated genotype calling after visual inspection, outlier exclusion, and manual adjustment of C_T_ threshold settings. Data were exported using an automated method available from the authors at request.

Samples with unequal calls across the TaqMan, PharmacoScan or Ion S5 CNV probes were subjected to L-PCR assays to generate products specific for *CYP2D6-2D7* or *CYP2D7-2D6* hybrids (amplicons E,G and H) as described,^9^ with minor modifications.

### The Agena MassARRAY

The Agena MassARRAY (Agena Bioscience, San Diego, CA) uses matrix-assisted laser desorption/ionisation-time of flight (MALDI-TOF) mass spectrometry technology for resolving oligonucleotides. We ran eight *CYP2C19* variants to enable calling of nine haplotypes. Genomic DNA was subjected to PCR followed by single-base extension with the extension products then being dispensed onto a SpectroCHIP Array and detected via mass spectrometry as described (Gaedigk et al., 2019)^28^ with haplotypes being assigned using Typer Analyzer software version v4.1.83 (Agena Bioscience).

## Results

### CYP2C19

There was a mean of 1112.19 mapped reads per *CYP2C19* variant in the Ion S5 data. Data for *CYP2C19* are shown in Table 1. The *CYP2C19* Ion S5 data were concordant with the prior genotypic information from the AmpliChip CYP450 and TaqMan *CYP2C19*17* genotyping, apart from two samples, in each of which a poor metabolizer variant not included in AmpliChip CYP450 Test but included in the Ion S5 AmpliSeq Pharmacogenomics Panel was detected. These adjusted the prior genotypes of *CYP2C19*1/*1* and *\*1/*2* to *\*1/*8* and *\*2/*6* respectively, with corresponding changes in enzyme phenotypic assignment from NM and IM to IM and PM respectively. The genotypes for these two samples by PharmacoScan and Agena were also *CYP2C19*1/*8* and*\*2/*6*. There were a couple of *CYP2C19*27* haplotypes (functionally indistinguishable from the *CYP2C19*1* and now classified as *CYP2C19*1.006*) in the PharmacoScan data^11^ For the *CYP2C19*2*, the Ion S5, PharmascoScan and TaqMan data were 100% concordant.

**Table 1.** Percentage concordance for Ion S5, PharmacoScan, and Agena with AmpliChip CYP450 and *CYP2C19*17*prior data by *CYP2C19* genotype.

| **AmpliChip and *CYP2C19*\*17 Genotype** | **N** | **Ion S5**<br>**(% concordance**<br>**of available data)** | **PharmacoScan**<br>**(% concordance**<br>**of available data)** | **Agena**<br>**(% concordance of**<br>**available data)** |
| --- | --- | --- | --- | --- |
| *\*1/\*1* | 44 | 95.4<sup>1</sup> | 97.7<sup>2</sup> | 95.4<sup>3</sup> |
| *\*1/\*17* | 24 | 100 | 100 | 100 |
| *\*1/\*2* | 17 | 94.1<sup>4</sup> | 94.1<sup>4</sup> | 94.1<sup>4</sup> |
| *\*17/\*17* | 1 | 0<sup>5</sup> |  | 100 |
| *\*2/\*17* | 4 | 100<sup>6</sup> | 100 | 100 |
| *\*2/\*2* | 2 | 100<sup>7</sup> | 100 | 100 |

<sup>1</sup>All *\*1/\*1* except one *\*1/\*8*, one no call
<sup>2</sup>All *\*1/\*1* except three (*\*1/\*27*, *\*1/\*8*, *\*17/\*27*; note that *CYP2C19*\*27 has been reclassified in PharmVar as *CYP2C19*\*1.006)
<sup>3</sup>One *\*1/\*8*, one no call
<sup>4</sup>All *\*1/\*2* except one *\*2/\*6*
<sup>5</sup>One no call
<sup>6</sup>All *\*2/\*17* with one *\*2.002/\*17*
<sup>7</sup>One *\*2/\*2.002*
For enhanced validation, another sample of *CYP2C19*\*17/\*17 genotype by AmpliChip was genotyped on IonS5 and PharmacoScan, and was *CYP2C19*\*17/\*17 by both technologies.

### CYP2D6

While the Ion S5 SNV data were of high quality (mean 1,111.13 mapped reads per variant), the CNV data had some no calls, some of which were rescued by manual calling. The CNV data from all platforms, where available, were used as a basis for grouping samples into *CYP2D6* gene copy number 1, 2, and 3, or hybrid pattern for further analysis.

Comparative genotypic and gene copy number data across the three platforms for samples with a CNV call of 1 are shown in Table 2. PharmacoScan and Ion S5 genotypes were consistent with the CNV probe data, which were also consistent across TaqMan, PharmacoScan and Ion S5 CNV probes. The AmpliChip genotypes were not consistent with the CNV probe data and therefore likely represent erroneous prior calls in 42.9% (6/14).

**Table 2.** Comparative *CYP2D6* data for samples with a CNV call of 1, with pairwise interplatform concordance (AmpliChip data requiring revisions resulting in adjustments in red)

| TaqMan CNV data |  |  | PScan CNV data |  |  | IonS5 | IonS5 | AmpliChip | PScan | IonS5 | AmpliChip vs PScan | AmpliChip vs IonS5 | PScan vs IonS5 |
| --- | --- | --- | --- | --- | --- | --- | --- | --- | --- | --- | --- | --- | --- |
| I2 | I6 | E9 | 5' | E9 | 3' | gene | E9 |  |  |  |  |  |  |
| 1 | 1 | 1 | 1 | 1 | 1 | 1 | 1 | *4XN/\*5* | *\*4/\*5* | *\*4/\*5* | No | No | Yes |
| 1 | 1 | 1 | 1 | 1 | 1 | 1 | 1 | *\*5/\*35* | *\*5/\*35* | *\*5/\*35* | Yes | Yes | Yes |
| 1 | 1 | 1 | 1 | 1 | 1 | 1 | 1 | *\*2/\*2* | *\*2/\*5* | *\*2.001/\*5* | No | No | Yes |
| 1 | 1 | 1 | 1 | 1 | 1 | 1 | 1 | *\*5/\*10* | *\*5/\*10* | *\*5/\*10* | Yes | Yes | Yes |
| 1 | 1 | 1 | 1 | 1 | 1 | 1 | 1 | *\*1/\*4* | *\*1/\*5* | *\*1/\*5* | No | No | Yes |
| 1 | 1 | 1 | 1 | 1 | 1 | 1 | 1 | *\*5/\*41* | *\*5/\*41* | *\*5/\*41* | Yes | Yes | Yes |
| 1 | 1 | 1 | 1 | 1 | 1 | 1 | 1 | *\*1/\*2XN* | *\*1/\*5* | *\*1/\*5* | No | No | Yes |
| 1 | 1 | 1 | 1 | 1 | 1 | 1 | 1 | *\*2/\*4* | *\*2/\*5* | *\*2.001/\*5* | No | No | Yes |
| 1 | 1 | 1 | 1 | 1 | 1 | 1 | 1 | *\*3/\*5* | *\*3.001/\*5* | *\*3/\*5* | Yes | Yes | Yes |
| 1 | 1 | 1 | 1 | 1 | 1 | 1 | 1 | *\*5/\*35* | *\*5/\*35* | *\*5/\*35* | Yes | Yes | Yes |
| 1 | 1 | 1 | 1 | 1 | 1 | 1 | 1 | *\*4/\*35* | *\*3.001/\*5* | *\*3/\*5* | No | No | Yes |
| 1 | 1 | 1 | 1 | 1 | 1 | 1 | 1 | *\*5/\*41* | *\*5/\*41* | *\*5/\*41* | Yes | Yes | Yes |
| 1 | 1 | 1 |  |  |  |  |  | *\*4/\*5* |  |  |  |  |  |

PScan = PharmacoScan, I2 = intron 2, I6 = intron 6, E9 = exon 9, 5' = 5' flanking region, 3' = 3' flanking region; vs = versus

For samples with a CNV call of two across the TaqMan or Pharmacoscan CNV probes, the concordance between Ion S5 and PharmacoScan was 94.4% (34/36), with the two non-concordant samples having variants detected by PharmacoScan that are not currently in the Ion S5 panel (*CYP2D6*22* and *CYP2D6*25*) (Supplementary Table 1). Samples with a CNV call of three consistently across probes for at least one platform were all genotyped as having a *CYP2D6XN* (i.e., duplication or multiplication of the *CYP2D6* gene) by at least one platform (Table 3). The AmpliChip-PharmacoScan concordance was higher than that for AmpliChip-IonS5 owing to the no calls in the Ion S5 data. While the AmpliChip provided haplotyping of the *CYP2D6XN* (i.e., assignment to one or other of the two chromosomes), neither PharmacoScan nor Ion S5 attempt to do that. *CYP2D6XN* positive samples comprised samples with the following haplotypes: *CYP2D6*1, *2, *3, *4, *6, *35*, and *\*41*. Haplotype assignment may be conducted by L-PCR followed by product clean-up and the use of relevant TaqMan assays.

**Table 3.**
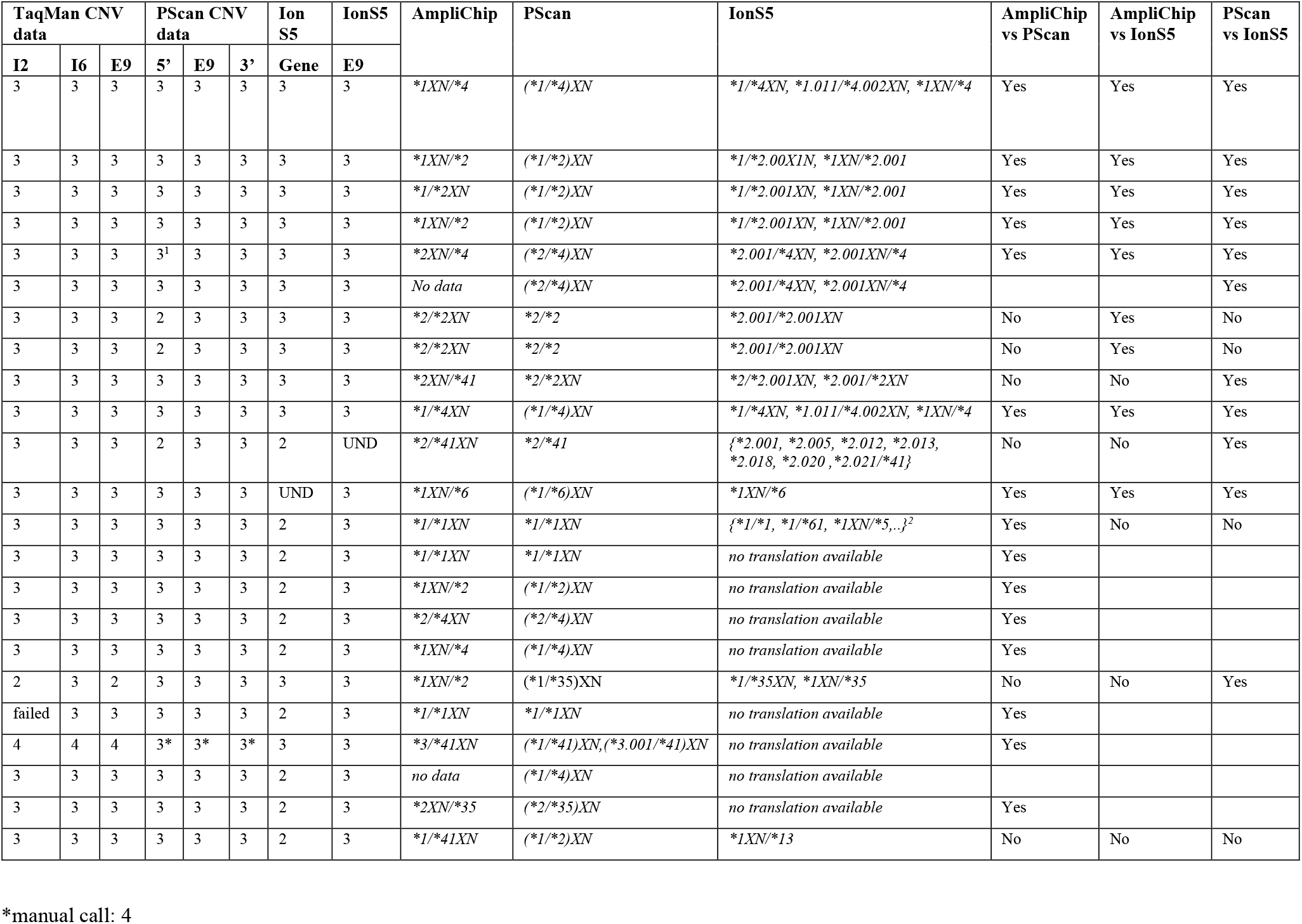
Comparative *CYP2D6* data for samples with a CNV call of 3, with pairwise inter-platform concordance.

| TaqMan CNV data |  |  | PScan CNV data |  |  | Ion S5 | IonS5 | AmpliChip | PScan | IonS5 | AmpliChip vs PScan | AmpliChip vs IonS5 | PScan vs IonS5 |
| --- | --- | --- | --- | --- | --- | --- | --- | --- | --- | --- | --- | --- | --- |
| I2 | I6 | E9 | 5' | E9 | 3' | Gene | E9 |  |  |  |  |  |  |
| 3 | 3 | 3 | 3 | 3 | 3 | 3 | 3 | *1XN/*4 | (*1/*4)XN | *1/*4XN, *1.011/*4.002XN, *1XN/*4 | Yes | Yes | Yes |
| 3 | 3 | 3 | 3 | 3 | 3 | 3 | 3 | *1XN/*2 | (*1/*2)XN | *1/*2.00X1N, *1XN/*2.001 | Yes | Yes | Yes |
| 3 | 3 | 3 | 3 | 3 | 3 | 3 | 3 | *1/*2XN | (*1/*2)XN | *1/*2.001XN, *1XN/*2.001 | Yes | Yes | Yes |
| 3 | 3 | 3 | 3 | 3 | 3 | 3 | 3 | *1XN/*2 | (*1/*2)XN | *1/*2.001XN, *1XN/*2.001 | Yes | Yes | Yes |
| 3 | 3 | 3 | 3<sup>1</sup> | 3 | 3 | 3 | 3 | *2XN/*4 | (*2/*4)XN | *2.001/*4XN, *2.001XN/*4 | Yes | Yes | Yes |
| 3 | 3 | 3 | 3 | 3 | 3 | 3 | 3 | No data | (*2/*4)XN | *2.001/*4XN, *2.001XN/*4 |  |  | Yes |
| 3 | 3 | 3 | 2 | 3 | 3 | 3 | 3 | *2/*2XN | *2/*2 | *2.001/*2.001XN | No | Yes | No |
| 3 | 3 | 3 | 2 | 3 | 3 | 3 | 3 | *2/*2XN | *2/*2 | *2.001/*2.001XN | No | Yes | No |
| 3 | 3 | 3 | 3 | 3 | 3 | 3 | 3 | *2XN/*41 | *2/*2XN | *2/*2.001XN, *2.001/*2XN | No | No | Yes |
| 3 | 3 | 3 | 3 | 3 | 3 | 3 | 3 | *1/*4XN | (*1/*4)XN | *1/*4XN, *1.011/*4.002XN, *1XN/*4 | Yes | Yes | Yes |
| 3 | 3 | 3 | 2 | 3 | 3 | 2 | UND | *2/*41XN | *2/*41 | {*2.001, *2.005, *2.012, *2.013, *2.018, *2.020, *2.021/*41} | No | No | Yes |
| 3 | 3 | 3 | 3 | 3 | 3 | UND | 3 | *1XN/*6 | (*1/*6)XN | *1XN/*6 | Yes | Yes | Yes |
| 3 | 3 | 3 | 3 | 3 | 3 | 2 | 3 | *1/*1XN | *1/*1XN | {*1/*1, *1/*61, *1XN/*5,...}<sup>2</sup> | Yes | No | No |
| 3 | 3 | 3 | 3 | 3 | 3 | 2 | 3 | *1/*1XN | *1/*1XN | no translation available | Yes |  |  |
| 3 | 3 | 3 | 3 | 3 | 3 | 2 | 3 | *1XN/*2 | (*1/*2)XN | no translation available | Yes |  |  |
| 3 | 3 | 3 | 3 | 3 | 3 | 2 | 3 | *2/*4XN | (*2/*4)XN | no translation available | Yes |  |  |
| 3 | 3 | 3 | 3 | 3 | 3 | 2 | 3 | *1XN/*4 | (*1/*4)XN | no translation available | Yes |  |  |
| 2 | 3 | 2 | 3 | 3 | 3 | 3 | 3 | *1XN/*2 | (*1/*35)XN | *1/*35XN, *1XN/*35 | No | No | Yes |
| failed | 3 | 3 | 3 | 3 | 3 | 2 | 3 | *1/*1XN | *1/*1XN | no translation available | Yes |  |  |
| 4 | 4 | 4 | 3* | 3* | 3* | 3 | 3 | *3/*41XN | (*1/*41)XN, (*3.001/*41)XN | no translation available | Yes |  |  |
| 3 | 3 | 3 | 3 | 3 | 3 | 2 | 3 | no data | (*1/*4)XN | no translation available |  |  |  |
| 3 | 3 | 3 | 3 | 3 | 3 | 2 | 3 | *2XN/*35 | (*2/*35)XN | no translation available | Yes |  |  |
| 3 | 3 | 3 | 3 | 3 | 3 | 2 | 3 | *1/*41XN | (*1/*2)XN | *1XN/*13 | No | No | No |

\*manual call: 4

Fifteen samples had an unequal call across at least two out of three CNV probesets (TaqMan or PharmacoScan or Ion S5); for seven of these, the CNV pattern was consistent with a *CYP2D7-2D6* hybrid, and for eight with a *CYP2D6-2D7* hybrid. For all of the *CYP2D6-2D7* hybrids, the pattern was consistent with an extra *CYP2D6* gene, either on the same haplotype as the hybrid gene (in “cis”), or on the other chromosome 22 (in “trans”). This was also the case for three of the *CYP2D7-2D6* hybrids. The expected L-PCR amplicons were generated for all of these 15 samples. Six samples had an unequal call across CNV probes for only one platform; four of these were genotyped as *CYP2D6* duplications (*CYP2D6*1X2/*4, *1X2/*1, *2X2/*1* and *\*2X2/*35*) and two as heterozygotes (*\*1/*2* and *\*1/*3.001*). For three of these, amplicon G was generated; however, it should be noted that the primer pair for this amplicon will also amplify up *CYP2D7* (Black et al., 2012,^9^ Figure 2, observed where the *CYP2D6* downstream gene was *\*1*, *\*4*, or *\*41;* these three all had genotypes including the *\*1* and/or *\*4*, specifically *\*1/*2, *1X2/*4, *4X2/*1*). Genotypes for the hybrids were predicted using the combination of available data and the samples may be further analysed by subjecting the L-PCR amplicons to genotyping.

## Discussion

For *CYP2C19*, the Ion S5-PharmacoScan genotype concordance was 100% for available data (91/91 samples with data on both platforms). Moreover, the Ion S5 and PharmacoScan *CYP2C19*2* data were 100% concordant with that from the TaqMan *CYP2C19*2* assay, and there were no adjustments required to the *CYP2C19*2* AmpliChip data. There were also no adjustments required to the prior *CYP2C19*17* data generated by TaqMan.

For *CYP2D6*, the IonS5 data showed a high rate of concordance with PharmacoScan for *CYP2D6* variants over a range of haplotypes tested apart from the *XN* and hybrid variants. Owing to the greater haplotype coverage by the PharmacoScan relative to the Ion S5, there were two samples with haplotypes identified only by the former (*CYP2D6*22* and *\*25*). To date it has been thought that it was not possible to employ next generation sequencing technology on genomic DNA for genotyping *CYP2D6;* rather, that one would have to first amplify up the whole gene and its surrounding regions and then use such technology.^28^ The Ion S5 AmpliSeq Pharmacogenomics Panel is to our knowledge the first product that uses a next generation platform to genotype *CYP2D6* variants from genomic DNA. Sixty-two different *CYP2D6* genotypes according to the AmpliChip CYP450 data were represented in the 96 samples analyzed herein, equating to 82.7% (65/75) of the range of AmpliChip genotypes in the GENDEP data. Haplotypes not represented in the prior AmpliChip data include the hybrid variants and others that are infrequent in Caucasians.^19,29^ We have herein demonstrated the feasibility of using this technology in this manner for a substantial proportion of the variety of *CYP2D6* genotypes and all of the common *CYP2C19* genotypes seen in Whites. We have also developed tools to facilitate the bioinformatic interpretation of such data into haplotypes of known function, so that enzyme activity can be deduced. Moreover, these tools could be adapted for the interpretation of data from specific variants in these genes arising from any other platform.

Both platforms were also able to call *CYP2D6*5* (gene deletion) variants in a manner consistent with each other and more reliably than the AmpliChip. Given the failure to detect *CYP2D6*5* reliably by the AmpliChip, the rest of available GENDEP samples were screened for *CYP2D6*5* using the TaqMan intron 2 and exon 9 probes, which revealed seven samples (7/853, 0.8%) with a CNV one call consistent with the presence of a previously undetected *CYP2D6*5* haplotype.

For the CNV call three samples (consistent with a duplicated *CYP2D6* gene or *XN* call, specifically *X2*, on one chromosome), the genotype on at least one platform was consistent with an *XN* call, with phase assignment of the *XN* being possible by amplifying the duplicated gene and using the resultant amplicon as a template for TaqMan SNV genotyping. By TaqMan CNV analysis, we have identified an additional 10 *XNs* in GENDEP beyond those reported here, giving a total of 33 *XNs* (33/853) and a frequency of 3.8%, consistent with prior data.^19^

A focus of recent research on *CYP2D6* is the hybrid haplotypes.^9,30^ Using TaqMan *CYP2D6* CNV probes,^31^ in addition to the hybrids haplotypes reported in this paper, we have identified data patterns consistent with 11 more potential hybrid haplotypes, making a total of 26/853 (3%) GENDEP samples. These remain to be further explored. Where CNV call was unequal across platforms, the particular pattern of CNV call was used to infer types of hybrid haplotype configurations, and hence which L-PCR set of primers should be employed to generate templates for SNV genotyping. Our previous validation work found a 100% concordance between TaqMan and DMETPlus *CYP2D6*41* genotyping and between TaqMan and PharmacoScan rs5758550 genotyping.^25,32,33^

The Luminex *CYP2C19* xTAG v3 has been validated against the Motorola Life Sciences eSensor DNA detection system.^34^ From 2006 onwards the AmpliChip *CYP2C19* genotypes were problematic owing to the identification of a relatively common variant conferring increased enzyme activity (*CYP2C19*17*)^35^ that was not covered by the assay. As this test was approved for *in vitro* diagnostic use by both the U. S. Federal Drug Administration and the European Medicines Agency, it could not be modified without going through costly regulatory approval again, and it was hence removed from the market amidst growing concerns regarding its comprehensiveness and sensitivity, despite its strengths including the apparent ability to discriminate heterozygous *CYP2D6XN* calls.

Data arising from the Luminex CYP2D6 xTAG v3 have been compared with the AmpliChip CYP450 Test, as well as with other genotyping platforms including AutoGenomics INFINITI, ParagonDx, and LDT SNaPShot.^36^ *CYP2D6* TaqMan assays have been compared with data arising from mPCR-RETINA, Sanger sequencing, long-PCR for *CYP2D6*5*, and next-generation sequencing data available via the 1000 Genomes Project.^37^ In a recent publication, 28 the range of *CYP2D6* haplotypes available in reference samples was increased by using multiple technologies including PharmacoScan, Agena iPLEX CYP2D6 v1.1, TaqMan assays, L-PCR, digital droplet PCR, next-generation sequencing of *CYP2D6* amplicons, and long-read single-molecule real-time sequencing of the whole *CYP2D6* gene. However, the use of a next generation sequencing platform to genotype genomic DNA was not included in the above, and much of the data interpretation appears to have been done manually, without automated methodology. While software exists to call *CYP2D6* haplotypes from next generation full sequencing data^38^, the novel contribution described herein is the use of the IonS5 AmpliSeq Panel, which employs short sequence reads as a genotyping tool and our haplotype derivation files for the interpretation of *CYP2D6* and *CYP2C19* data arising.

We propose the following methodology for genotyping *CYP2D6:* use a high throughput platform (e.g., Ion S5, PharmacoScan or Agena) with automated haplotype translation plus TaqMan *CYP2D6* CNV analysis, followed by L-PCR for gene duplications or hybrids and using the resultant amplicons as templates for genotyping (potentially adapting the same high throughput platform as used for genotyping the genomic DNA), and compare the data from the amplicons with that from genomic DNA to derive genotypes for the relevant *CYP2D6* structural variants. The utilization of a greater number of *CYP2D6* CNV probes by the platforms where necessary and improvements in performance of such probes where indicated should enhance the feasibility of this suggested workflow.

In addition to the complexity of the locus, another barrier that has limited *CYP2D6* to date has been the cost of the genotyping technology. The cheapest way to genotype is in fact by sequencing. The most high throughput (and therefore potentially most cost-effective) sequencing methodology is NGS. And while we have developed haplotype derivation files specifically for use with the Ion S5, they could be adapted for other NGS platforms such as Illumina. Should an NGS provider develop a panel including not only a greater number of *CYP2D6* SNVs covered than those included in the Ion S5 AmpliSeq Panel but also a greater number of more reliable *CYP2D6* CNV probes and offer this at a competitive price, this could be a game-changer: moving *CYP2D6* from the position of being a locus that by virtue of its challenging complexity has been inconsistently genotyped to one for which genotyping becomes accessible and affordable.

## Conclusion

The inter-platform concordance for *CYP2C19* was high (94-98%). For *CYP2D6*, the IonS5-PharmacoScan concordance was 94% for a range of variants tested apart from those with at least one extra copy of a *CYP2D6* gene or a hybrid. We have also demonstrated feasibility of using a NGS platform for genotyping *CYP2D6* and *CYP2C19*, and, moreover, that this provides adequate coverage of *CYP2D6* for approximately 94% of Whites. Further, we have generated automated methods for translating NGS *CYP2D6* sequencing data into *CYP2D6* haplotypes. We suggest a minimum set of assays for *CYP2D6* and *CYP2C19* as follows: a multiplex platform in combination with specific TaqMan probes for detecting *CYP2D6* copy number, with further analysis of a subset of such data as necessary. *CYP2D6* and *CYP2C19* genotyping may thereby become much more readily accessible, with potential substantial savings in health care costs and public benefit.

## Acknowledgments

We thank Lin Chen for support with interpretation of data arising from the AmpliSeq Pharmacogenomics Panel, and Carsten A. Bruckner for support for PharmacoScan data interpretation including supplying the manufacturer’s version of the *CYP2D6* translation file. The work reported herein was funded by: a Canada Foundation for Innovation (CFI), John R. Evans Leaders Fund (JELF) grant (32147 - Pharmacogenetic translational biomarker discovery), an Alberta Innovates Strategic Research Project (SRP51_PRIME - Pharmacogenomics for the Prevention of Adverse Drug Reactions in mental health; G2018000868), an Alberta Centennial Addiction and Mental Health Research Chair (to KJA), Alberta Innovation and Advanced Education Small Equipment Grants Program (to KJA), the Neuroscience and Mental Health Institute, Department of Psychiatry, and the Faculty of Medicine and Dentistry at the University of Alberta. Infrastructure used in this project from GP’s lab was supported by a Hotchkiss Brain Institute Dementia Equipment Fund grant (to CB and GP for the Ion S5) and a Canada Foundation for Innovation John R Evans Leaders Fund Grant (CFI-JELF) (36624 - Neuromuscular genetics program, to GP). GENDEP was funded by a European Commission Framework 6 grant, LSHB-CT-2003-503428. Roche Molecular Systems supplied the AmpliChip CYP450 Test arrays and some associated support. GlaxoSmithKline and the Medical Research Council (UK) contributed by funding add-on projects in the London centre. This paper represents independent research part-funded by the National Institute for Health Research (NIHR) Biomedical Research Centre at South London and Maudsley NHS Foundation Trust and King’s College London. The views expressed are those of the authors and not necessarily those of the NHS, the NIHR or the Department of Health and Social Care.

## Notes

### Competing Interest Statement

N.H. has participated in research supported by CSF project No. IP-09-2014-2979. D.S. has received grant/research support from Janssen and Lundbeck, and has served as a consultant or on advisory boards for Janssen and Lundbeck. C.A.B. is a member of the Clinical Pharmacogenetics Implementation Consortium and the Pharmacogene Variation Consortium; he has also participated in collaborative research with employees of MyDNA (a pharmacogenetic testing company) but does not have equity, stocks, or options in this company or any other pharmacogenetic companies. K.J.A. is a member of the Clinical Pharmacogenetics Implementation Consortium and the Pharmacogene Variation Consortium, has received two research grants in the last two years from Janssen Inc., Canada (fellowship grants for trainees) and provided consultancy services (unpaid) for HLS Therapeutics. All other authors have nothing to disclose.

### Summary of Updates

Updated authors, disclosures, and updates to the paper including added data to tables and supplemental files.

